# Resting-state network topology and planning ability in healthy adults

**DOI:** 10.1101/696856

**Authors:** Chris Vriend, Margot J. Wagenmakers, Odile A. van den Heuvel, Ysbrand D. van der Werf

## Abstract

Functional magnetic resonance imaging (fMRI) studies have been used extensively to investigate the brain areas that are recruited during the Tower of London (ToL) task. Nevertheless, little research has been devoted to study the neural correlates of the ToL task using a network approach. Here we investigated the association between functional connectivity and network topology during resting-state fMRI and ToL task performance, that was performed outside the scanner. Sixty-two (62) healthy subjects (21-74 years) underwent eyes-closed rsfMRI and performed the task on a laptop. We studied global (whole-brain) and within subnetwork resting-state topology as well as functional connectivity between subnetworks, with a focus on the default-mode, fronto-parietal and dorsal and ventral attention networks. Efficiency and clustering coefficient were calculated to measure network integration and segregation, respectively, at both the global and subnetwork level. Our main finding was that higher global efficiency was associated with slower performance (β = .22, P_bca_ = .04) and this association seemed mainly driven by inter-individual differences in default-mode network connectivity. The reported results were independent from age, sex, education-level and motion. Although this finding is contrary to earlier findings on general cognition, we tentatively hypothesize that the reported association may indicate that individuals with a more integrated brain during the resting-state are less able to further increase network efficiency when transitioning from a rest to task state, leading to slower responses. This study also adds to a growing body of literature supporting a central role for the default-mode network in individual differences in cognitive performance.

## Introduction

Executive functions are a set of mental processes that enable us to plan, focus attention, remember instructions and handle several tasks at once (Diamond 2013). Functional magnetic resonance imaging (fMRI) studies have shown that these functions are associated with functional connectivity (FC) of certain resting-state networks (RSN) (Rabinovici et al. 2015; Funahashi and Andreau 2013; Nowrangi et al. 2014). Various RSN have been shown to be involved in executive functions, including the default mode network (DMN) which is active during rest and deactivates during task performance (Buckner et al. 2008; Mak et al. 2017; Anticevic et al. 2012). Other relevant RSN for cognition are the frontoparietal network (FPN) (Cole et al. 2014b; Cole et al. 2012) and the dorsal and ventral attention networks (DAN and VAN, respectively) (Fortenbaugh et al. 2018). Although the utility of RSN in cognitive neuroscience and understanding of the neural correlates of cognition has been debated (Campbell and Schacter 2017; Davis et al. 2017; Iordan and Reuter-Lorenz 2017), resting-state FC patterns show good correspondence with task-based FC patterns (Krienen et al. 2014), are fundamentally stable (Gratton et al. 2018) and may act as an intrinsic network architecture that shapes FC when evoked by a cognitive task (Cole et al. 2014a; Ito et al. 2017).

The architecture or topology of the brain can be studied using graph analysis, where the brain is simplified to a graph of nodes (i.e., different brain regions) and edges (i.e., connections between brain regions) (Wang et al. 2010; Bullmore and Sporns 2009). Different properties of the brain network can be calculated using this graph. For example, efficiency and clustering describe the ability of a network to integrate and segregate information, respectively (Cohen and D’Esposito 2016; Lord et al. 2017). The brain balances its ability to integrate and easily transmit information throughout the network, and to segregate information processing in clusters of highly interconnected (specialized) neighboring nodes (Bullmore and Sporns 2009). This ability of the brain for integration and segregation is vital for cognitive processes (Cohen and D’Esposito 2016) and higher intelligence has been associated with a more efficient network topology (Langer et al. 2012; van den Heuvel et al. 2009). Conversely, dementia and cognitive impairments in the light of brain disorders generally show dysfunction in the brain’s ability to functionally integrate and segregate information (Dai et al. 2019; Lopes et al. 2017; Rocca et al. 2016). Nevertheless, studies on the associations between network topology and inter-individual differences in cognitive functions in healthy subjects are relatively scarce, e.g. (Cohen and D’Esposito 2016; Sheffield et al. 2017), and to the best of our knowledge, no study has yet focused on the association between network topology and planning capacity. Planning is the ability to think ahead in order to achieve a goal via a series of intermediate steps (Owen 1997) and is a vital function in daily life that we here operationalize in the form of the Tower of London (ToL) task. In this study, we investigated the association between RSN topology and planning performance, using a graph-based approach. Based on prior research (Langer et al. 2012; van den Heuvel et al. 2009; Sheffield et al. 2017), we hypothesized a positive relationship between network topology measured during resting-state and cognitive planning ability, measured using the ToL task performed outside of the scanner.

## Methods

### Subjects and measurements

Data of healthy adult controls from two previous case-control studies (Gerrits et al. 2015; de Wit et al. 2012) were pooled for the current study. Exclusion criteria for all healthy subjects were the use of psychoactive medication, current or past psychiatric diagnosis, a history of a major physical or neurological illness, MRI contraindications or a history of alcohol abuse. Further exclusion criteria for the current study were: no available data on the ToL task, extreme behavioral scores (≥ 2 SD from the mean), a time-interval of more than 21 days between resting-state fMRI (rs-fMRI) and performing the ToL task, or pathological incidental findings on the structural MRI scan. Written informed consent was provided by all participants according to the Declaration of Helsinki and the studies were approved by the Medical Ethical Committee of the VU University Medical Centre (Amsterdam, The Netherlands).

The participants performed a computerized version of the ToL task as a measure of planning (Phillips et al. 2001; Shallice 1982). Details of the ToL task are provided in the study by (van den Heuvel et al. 2003). In short, the participants saw two configurations (“begin” and “goal” position) of three colored beads on vertical posts of different heights. The purpose of the task is to determine the minimum number of moves (1, 2, 3, 4, or 5) needed to match the configuration of the goal position. Participants responded via the matching keyboard-button. The first post can hold all three beads, the second two, and the third post one. Only one bead can be moved at a time and only if there is no other bead on top of it. Prior to the experiment, participants were provided verbal and written explanation and performed a practice run. Performance on the ToL task was indicated by the mean accuracy and mean reaction time on correct trials across all five difficulty levels (Kaller et al. 2016). Intelligence scores were approximated by the Dutch Adult Reading test (NLV; (Schmand et al. 1991). We scored education level according to the Dutch Verhage scale (Verhage 1964) that ranges from 1 - *primary school not finished*, to 7 – *university or higher.* Handedness was assessed using the Edinburgh Handedness Inventory (Oldfield 1971).

### MR Image acquisition

MR images were acquired at Amsterdam UMC, location VUmc (Amsterdam, The Netherlands) on a GE Signa HDxt 3 Tesla MRI scanner (General Electric, Milwaukee, WI) with an eight channel head coil. The participant’s head was immobilized using foam pads to reduce motion artifacts. Participants were told to lie still, keep their eyes closed and not fall asleep during the acquisition of the rs-fMRI scan (duration: 5.9 min). T2*-weighted echo-planar (EPI) images were acquired with TR = 1.8 sec, TE = 35 ms, 64×64 matrix, field of view = 24 cm and flip angle = 80° and 40 ascending slices per volume (3.75 x 3.75 mm in plane resolution; slice thickness = 2.8 mm; interslice gap = 0.2 mm). Structural scanning encompassed a sagittal three-dimensional gradient-echo T1-weighted sequence (256 x 256 matrix; voxel size = 1 x 0.977 x 0.977 mm; 172 slices).

### Image (pre)processing

RS-fMRI and T1-weighted images were preprocessed with FMRIB’s Software Library version 5.0.10 (FSL; (Smith et al. 2004)). The first four volumes were discarded to reach steady-state magnetization. Non-brain tissue was removed using BET and the structural image was segmented into gray (GM), white matter (WM) and cerebrospinal fluid (CSF) using FAST. Functional images were re-aligned using McFLIRT and the resulting six rigid-body parameters were used to calculate the motion parameters. Functional images were spatially smoothed with a 5 mm full width at half maximum (FWHM) kernel. Subjects with significant motion during scanning, defined as a mean relative root mean squared displacement (RMS) > 0.2 mm, or > 20 volumes with frame-wise relative RMS displacement > 0.25 mm, were excluded (Ciric et al. 2017). Because rs-fMRI is exceptionally sensitive to motion artefacts (Power et al. 2015), we additionally performed ICA-AROMA (Pruim et al. 2015). ICA-AROMA is a single-subject denoising strategy based on independent component analysis (ICA) that automatically identifies motion-related components in the functional data based on their high-frequency content, correlation with the motion parameters and edge and CSF fraction and removes their variance from the data (Pruim et al. 2015). ICA-AROMA has been shown to provide a good trade-off between reducing noise and preserving BOLD signal (Ciric et al. 2017; Pruim et al. 2015; Parkes et al. 2018). After ICA-AROMA, additional nuisance regression was performed by removing signal from the WM and CSF and functional images were high-pass filtered (100 seconds cut-off).

The functional scan was registered to the anatomical T1-scans using boundary based registration (FSL epi_reg). The anatomical image was parcellated into 225 nodes; 210 cortical nodes were defined based on the Brainnetome Atlas (Fan et al. 2016), 14 subcortical nodes were individually segmented using FSL FIRST (Patenaude et al. 2011) and one cerebellar node was defined based on the FSL’s cerebellar atlas (Diedrichsen et al. 2009). EPI distortions during fMRI can lead to signal drop-out. To account for signal dropout near air/tissue boundaries during scanning, we applied a mask to the functional scan to exclude voxels with signal intensities in the lowest quartile of the robust range (Meijer et al. 2017). Nodes were discarded if they comprised less than four signal-containing voxels. This rendered a total of 194 common brain regions across all subjects. Time-series were extracted from each node. The cortical nodes were subdivided into four RSN: the DMN, FPN, DAN and VAN based on the functional subdivision by Yeo et al. (2011); see supplementary table 1.

### Functional connectivity matrices

To measure FC and construct connectivity matrices we applied wavelet coherence on the time-series of each possible pair of the 194 brain regions within the frequency range 0.06 and 0.12 Hz (Chang and Glover 2010). Wavelet coherence has several advantages over Pearson’s correlations, including denoising properties and robustness to outliers (Gu et al. 2017; Fadili and Bullmore 2004; Achard et al. 2006). The 0.06-0.12Hz frequency range was chosen because it has been suggested to be a reliable and robust range that is associated with cognitive performance (Zhang et al. 2016; Bassett et al. 2013). We applied wavelet coherence to the entire rs-fMRI scan to calculate the network measures (see below). An overview of the (pre)processing pipeline is provided in Figure 1.

**Figure 1.**
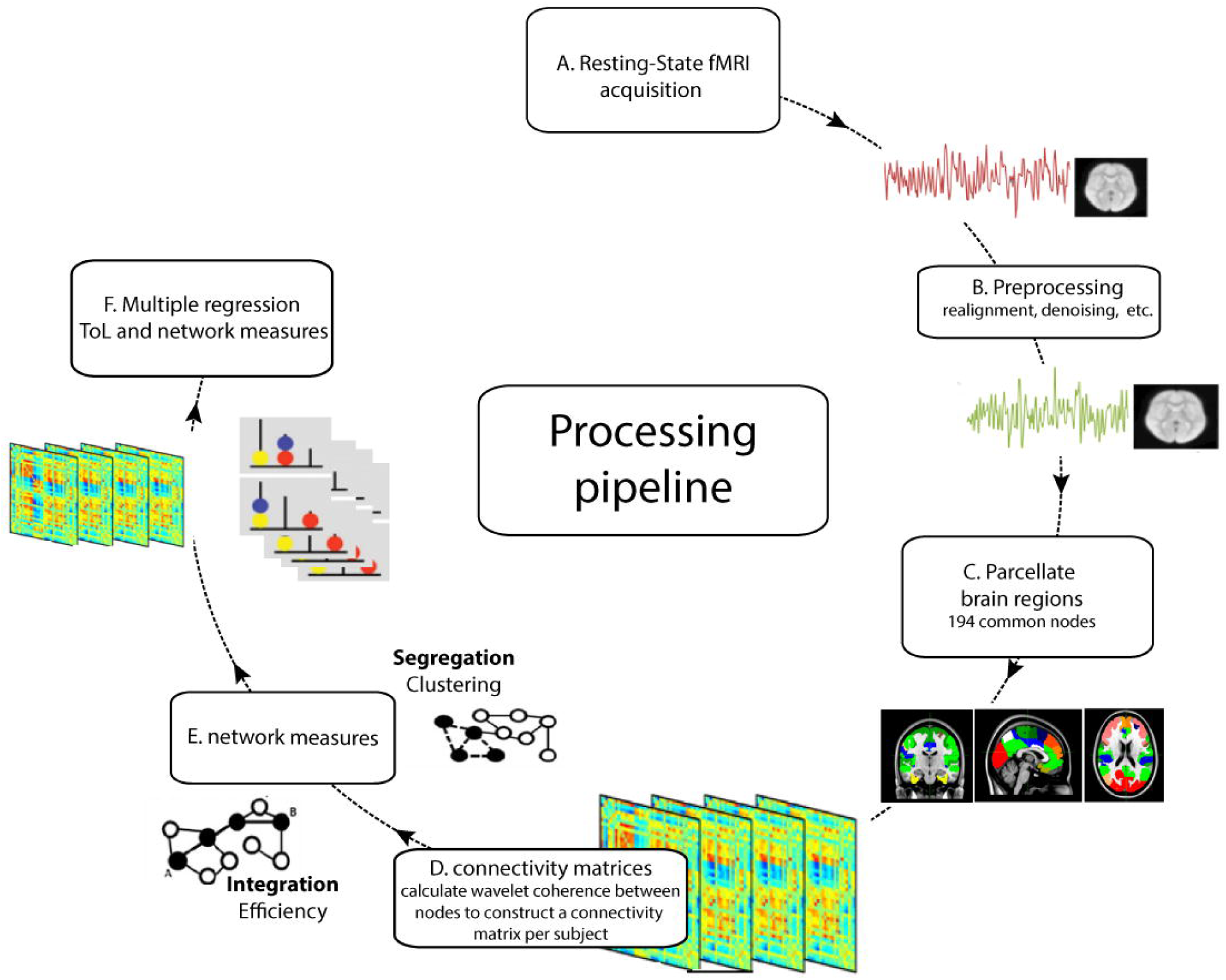
Outline of the processing pipeline. (A) Resting-state fMRI data were collected and (B) pre-processed. The brain was (C) parcellated into separate brain regions (nodes). There were 194 nodes common to all subjects with enough signal to (D) construct connectivity matrices (see text) using wavelet coherence. (E) network measures were calculated from each connectivity matrix on the global and subnetwork level. (F) multiple regression analyses were applied to relate performance on the Tower of London (ToL) task to network measures.

### Network measures

At the global level we calculated global efficiency and global clustering coefficient (Gcc). Global efficiency is the inverse of the average path length (i.e. the maximum connectivity between each pair of nodes), with high efficiency meaning that information can rapidly travel through the whole network (Latora and Marchiori 2001). Gcc is equivalent to the proportion of the actual number of edges between the nearest neighbors of a node to all possible edges and signifies the tendency of the whole network to segregate into locally interconnected triplets that function as a specialized subunit (Rubinov and Sporns 2010). Test-retest reliability of global efficiency and Gcc are fair-to-good (Welton et al. 2015). At the subnetwork level, we calculated efficiency and clustering coefficient for each of the four RSNs (DMN, FPN, DAN and VAN). In addition, we determined the mean FC *between* each of the four RSNs (resulting in six between-network mean FC values). Test-retest reliability of these measures at the subnetwork level is unknown.

### Data analysis

Statistical analyses were performed using SPSS version 25 (IBM Corp, Armonk, NY, USA). We describe demographical characteristics and performance on the ToL task using means and standard deviations, unless indicated otherwise. Pearson’s (r) or Spearman’s rho (r_s_) correlations were performed between demographic and performance measures, depending on the distribution. We performed bootstrapped hierarchal multiple regression analysis to investigate the association between network measures (predictors) and accuracy and reaction time on the ToL task (outcome measures). Because age was correlated with performance, age was entered in the first block of all models. The network measure of interest and mean RMS displacement, as a measure for motion, were entered in the second and third block, respectively. As a sensitivity analysis, we entered sex or education level to the fourth block of the model. The regression models were bootstrapped using 2000 iterations. We report bias and accelerated (BCa) confidence intervals and the accompanying P-values (P_bca_) as they account for bias and skewness in the data and provide a more robust estimate of the association that is less reliant on the distribution of the variable. All assumptions of multiple regression analyses, including homoscedasticity of residuals, were assessed and met. We performed separate analyses for the network measures on the global level and on the subnetwork level. On the subnetwork level, type I errors due to multiple comparisons were minimized using the False Discovery Rate (FDR, q<.05 (Benjamini and Hochberg 1995)). Statistical significance was set to p < 0.05 for all analyses. No formal power analysis was conducted prior to the execution of this study.

## Results

### Sample characteristics and behavioral results

Of the 69 participants with an available ToL task and rs-fMRI data, seven had to be excluded (see figure 2), which resulted in a total sample size of 62 participants, aged between 21 and 74 years old (M_age_ = 48.1 ± 13.9, 33 males). The time between performing the ToL task and the rs-fMRI was on average 6.2 ± 4.6 [range: 0-21] days. See Table 1 for the sample characteristics. Age showed a positive correlation with reaction time (*r* = .498, *p* < .001) but only a trend-level negative correlation with accuracy (*r* = -.243, *p* = .057) indicating that older participants tended to respond slower and slightly less accurately. The average motion during rs-fMRI (expressed as mean relative RMS framewise displacement) was 0.068 ± 0.029 [range: 0.027-0.17] and was positively correlated with age (*r*_*s*_ = .34, *p* = .007) but not performance on the ToL (reaction time: *r*_*s*_ = .06, *p* = .66; accuracy: *r*_*s*_ = -.18, *p* = .12).

**Table 1.** Sample characteristics.

|  |  |
| --- | --- |
| N subjects (% female) | 62 (46.8) |
| Age (years) | 48.1 ± 13.9 |
| Education level (in %)# |  |
| 3 | 1.6 |
| 4 | 48 |
| 5 | 29.0 |
| 6 | 43.5 |
| 7 | 17.7 |
| Handedness (R/L)* | 54/7 |
| ToL accuracy (%) | 87.7 ± 7.5 |
| ToL reaction time (s) | 10.1 ± 2.1 |
| Mean relative RMS | 0.07 ± 0.03 |

**Figure 2.**
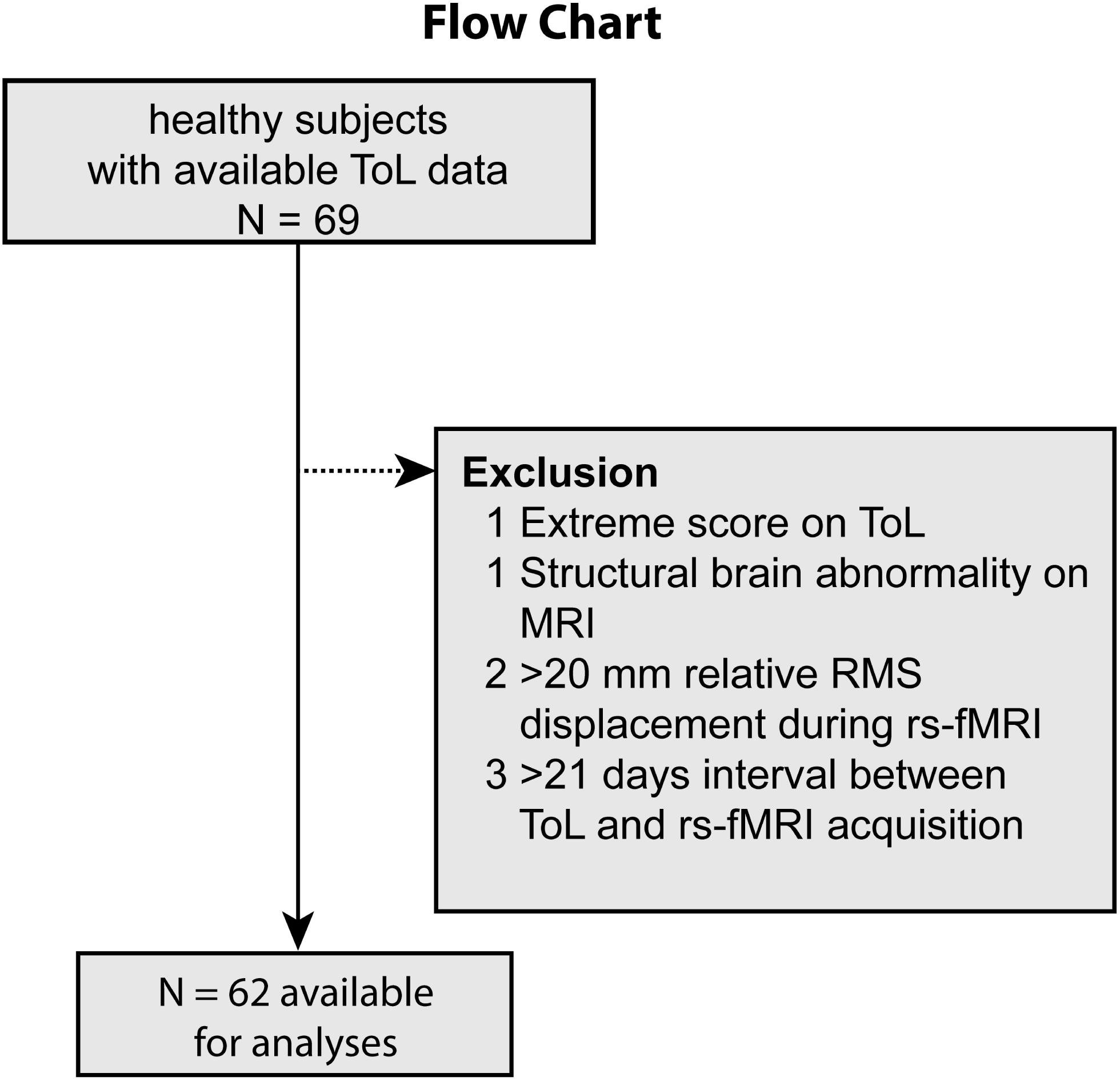
Flowchart of participant exclusion.

### Global topology

Global efficiency (β = .22, P_bca_ = .04) but not Gcc (β = -.09, P_bca_ = .57) was positively associated with reaction time above and beyond the effects of age (see Table 2). There were no significant associations with accuracy. Adding sex or education level as a nuisance covariate to the model had no effect on these results.

**Table 2.** associations between TOL performance and global network measures.

| TOL | Model | B (SE) | 95% CI (BCa) | Beta | P<sub>BCa</sub> | R<sup>2</sup> |
| --- | --- | --- | --- | --- | --- | --- |
| RT | Age | 0.09 (0.015) | 0.06, 0.12 | .595 | <.001 |  |
|  | GE | 9.79 (4.72) | 0.84, 19.8 | .219 | .039 | .293 |
|  | Motion | -16.46 (6.53) | -30.6, -5.7 | -.230 | .008 |  |
|  | Age | 0.08 (0.017) | 0.04, 0.11 | .531 | <.001 |  |
|  | Gcc | -32.88 (53.72) | -133.9, 68.8 | -.089 | .567 | .252 |
|  | Motion | -12.03 (7.69) | -26.6, 2.3 | -.168 | .113 |  |
| ACC | Age | -0.131 (.073) | -0.28, -0.009 | -.241 | .079 |  |
|  | GE | -31.1 (21.57) | -76.3, 9.6 | -.194 | .156 | .06 |
|  | Motion | -21.62 (29.23) | -69.0, 43.1 | -.084 | .450 |  |
|  | Age | -0.09 (.075) | -0.24, 0.05 | -.164 | .249 |  |
|  | Gcc | 206.28 (155.46) | -79.3, 515.3 | .154 | .187 | .05 |
|  | Motion | -41.39 (27.02) | -89.5, 13.3 | -.161 | .117 |  |

For each analysis, age was entered in model 1, the network measure in model 2 and motion parameters in model 3. Only the results of model 3 are shown here. P-values are bootstrapped using 2000 permutations. Abbreviations: TOL = Tower of London task, RT = reaction time, ACC = accuracy, SE = Standard Error, CI = confidence interval, BCa= Bias corrected and accelerated, GE = Global Efficiency, Gcc = Global Clustering Coefficient. Motion was defined as the mean root-mean-squared framewise displacement during the entire resting-state MRI scan.

### Subnetwork topology

Both efficiency and clustering of the DMN (efficiency: β = .25, P_bca_ = .018; clustering: β = .23, P_bca_ = .039) but not the other subnetworks (see supplemental Table 2) were positively related to reaction time. These associations did not, however, survive the multiple comparison correction (DMN efficiency P_fdr_ = 0.072; DMN clustering: P_fdr_ = 0.077). Adding sex or education level as additional nuisance covariate to the model had no effect on the results. Consistent with the results on the global level, there were no significant associations with accuracy of task performance (Supplemental Table 3).

### Between-subnetwork connectivity

FC between the DMN and FPN (β = .23, P_bca_ = .04), the DAN (β = .21 P_bca_ = .04) and the VAN (β = .20, P_bca_ = .04) were all positively associated with reaction time. These associations did not survive the FDR correction for multiple comparisons (all P_fdr_ = .09; supplemental Table 4 and 5).

### Post-hoc analyses

Because of possible floor/ceiling effects during the less demanding 1, 2 and 3 step trials of the ToL task, we re-ran the regression models using only the mean accuracy rates and reaction times during ToL steps 4 and 5. These post-hoc analyses showed that at the global level reaction time – but not accuracy – was still associated with global efficiency (β = .27, P_bca_ = .04), not Gcc (β = -.09, P_bca_ = .59). At the subnetwork level, efficiency of the DMN (β = .41, P_fdr_ *=* .001) and FPN (β = .32, P_fdr_ *=* .045), clustering of the DMN (β = .33, P_fdr_ *=* .05) and FC between the DMN and FPN (β = .38, P_fdr_ *=* .02) were all positively associated with reaction time, after FDR correction for multiple comparisons.

## Discussion

In this study in 62 healthy adults with a wide age range we investigated the association between network topology during a rs-fMRI session and cognitive planning ability during a ToL task that was performed outside the scanner. We observed that global (whole-brain) efficiency was associated with reduced planning speed and that this effect was mainly driven by the FC of the DMN. The results were independent from inter-individual differences in age, gender, education level and motion during rs-fMRI. Post-hoc analyses showed that our results were strongest when focusing on the higher task load trials of the ToL task (four and five step trials).

Global efficiency provides a measure of how well-integrated a network is and how easily information can travel from one node to another on the other side of the network, while the clustering coefficient is a measure of how well-connected nodes are locally into segregated triangles of neighboring nodes. Both measures are often used to describe the characteristics of a network and abnormalities in these network measures are commonly observed in the structural and functional networks of patients with a brain disorder (Bullmore and Sporns 2012; Griffa et al. 2013; Worbe 2015; Lord et al. 2017). Here we observed that subjects with a higher global efficiency show slower planning performance on the ToL task. This finding is at odds with our hypothesis and previous studies that observed that higher global efficiency is associated with higher global intelligence (van den Heuvel et al. 2009; Sheffield et al. 2017) and performance on working memory tasks (Cohen and D’Esposito 2016; Sheffield et al. 2015). One other study has also previously found that a higher global efficiency was associated with worse performance on a working memory task, but only in older adults and only when focusing on task-based FC (Stanley et al. 2015). This is the first study, however, to investigate planning ability. One possible, albeit less plausible, explanation might therefore be that planning requires a different whole-brain network organization than working memory tasks or general intelligence. Alternatively, the higher global efficiency in individuals with slower performance on the ToL task may also point towards a more random network (Ajilore et al. 2014). As there was no association between ToL task speed and lower global clustering (a characteristic feature of random networks), this explanation is also less viable.

Studies have shown that, although the resting-state provides a core and intrinsic network architecture that highly overlaps with the network topology of task-states (Cole et al. 2010; Krienen et al. 2014), significant reorganization does take place during the execution of tasks, and the magnitude and spatial redistribution depends on the task and its load (Cohen and D’Esposito 2016; Davison et al. 2015). Furthermore, the ease with which a network can reconfigure from rest to task-states correlates with task performance and general cognition (Braun et al. 2015; Bassett et al. 2011; Telesford et al. 2016; Hearne et al. 2017). Transitions of rest to (demanding) task-states have generally been associated with an increase in global efficiency, signifying a better integrated network (Cohen and D’Esposito 2016; Hearne et al. 2017; Shine et al. 2016; see Shine and Poldrack 2018 for a review). This increase in network integration is, however, not unconstrained, as a fully integrated functional network would lead to epileptic seizures and violates the principles of cost-efficiency (Shine and Poldrack 2018; Bullmore and Sporns 2012). Assuming that in our subjects network integration would similarly increase from the resting-state to task-state, i.e. execution of the ToL task, it is conceivable that global efficiency could not increase sufficiently in those subjects with an already highly integrated network during the resting-state to meet task demands, leading to a slower behavioral response. This concept is schematically depicted in Figure 4. Although this hypothesis receives indirect support from multiple previous studies on dynamic network reconfigurations (Shine and Poldrack 2018), we unfortunately did not acquire fMRI scans during execution of the ToL task and therefore this explanation currently remains speculative. Because the slower responses were not associated with lower accuracy (r_s_ = -.19, P = .13) and we did not observe an association between network topology and accuracy, our results may not be specific for planning performance but may also be related to an overall slower information processing speed. Why we did not find an association with task accuracy is currently unclear.

**Figure 3.**
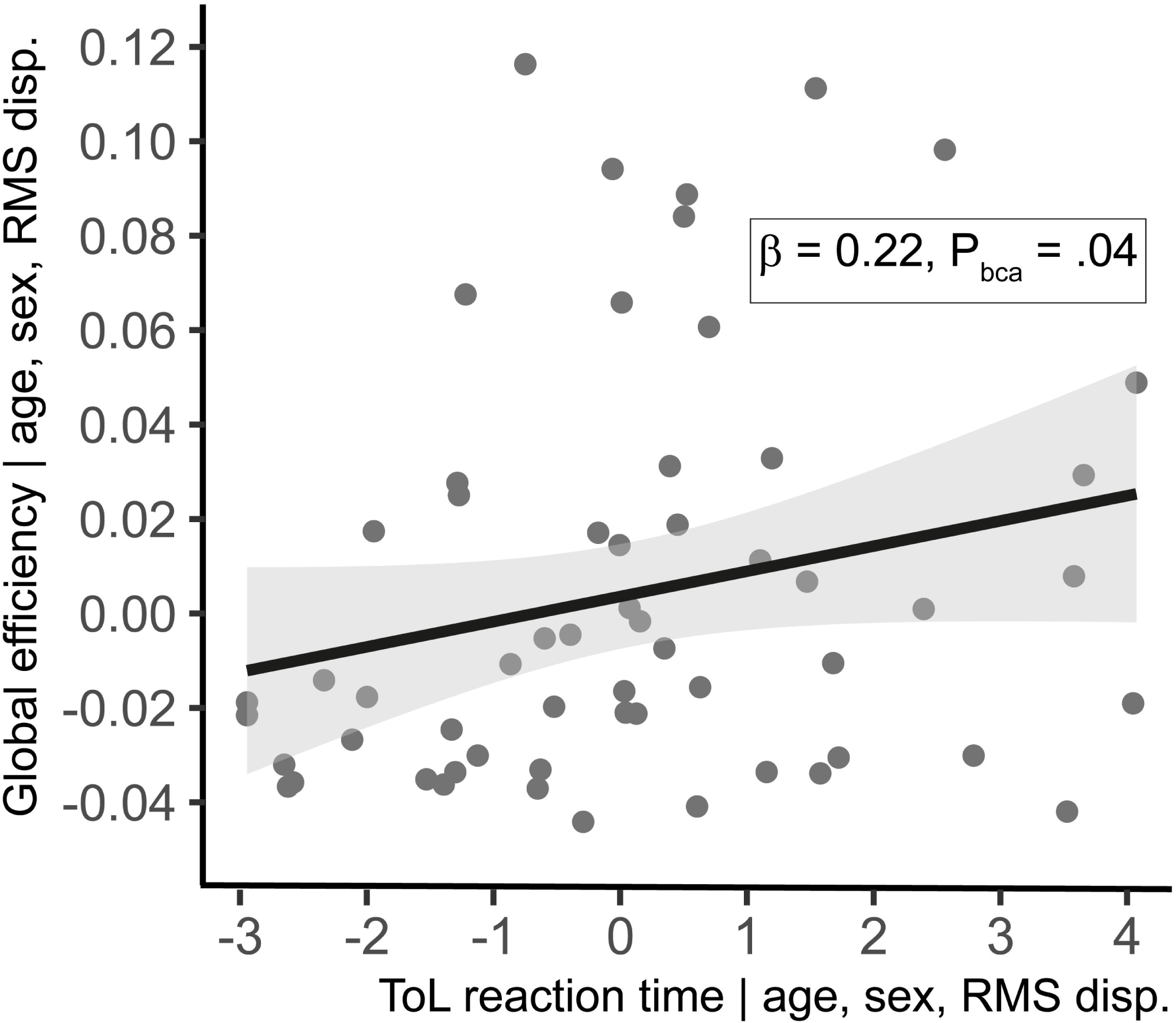
Partial correlation plot of association between reaction time on the Tower of London task and Global (whole-brain) efficiency. Abbreviations: ToL = Tower of London. RMS disp. = mean root-mean-squared framewise displacement.

**Figure 4.**
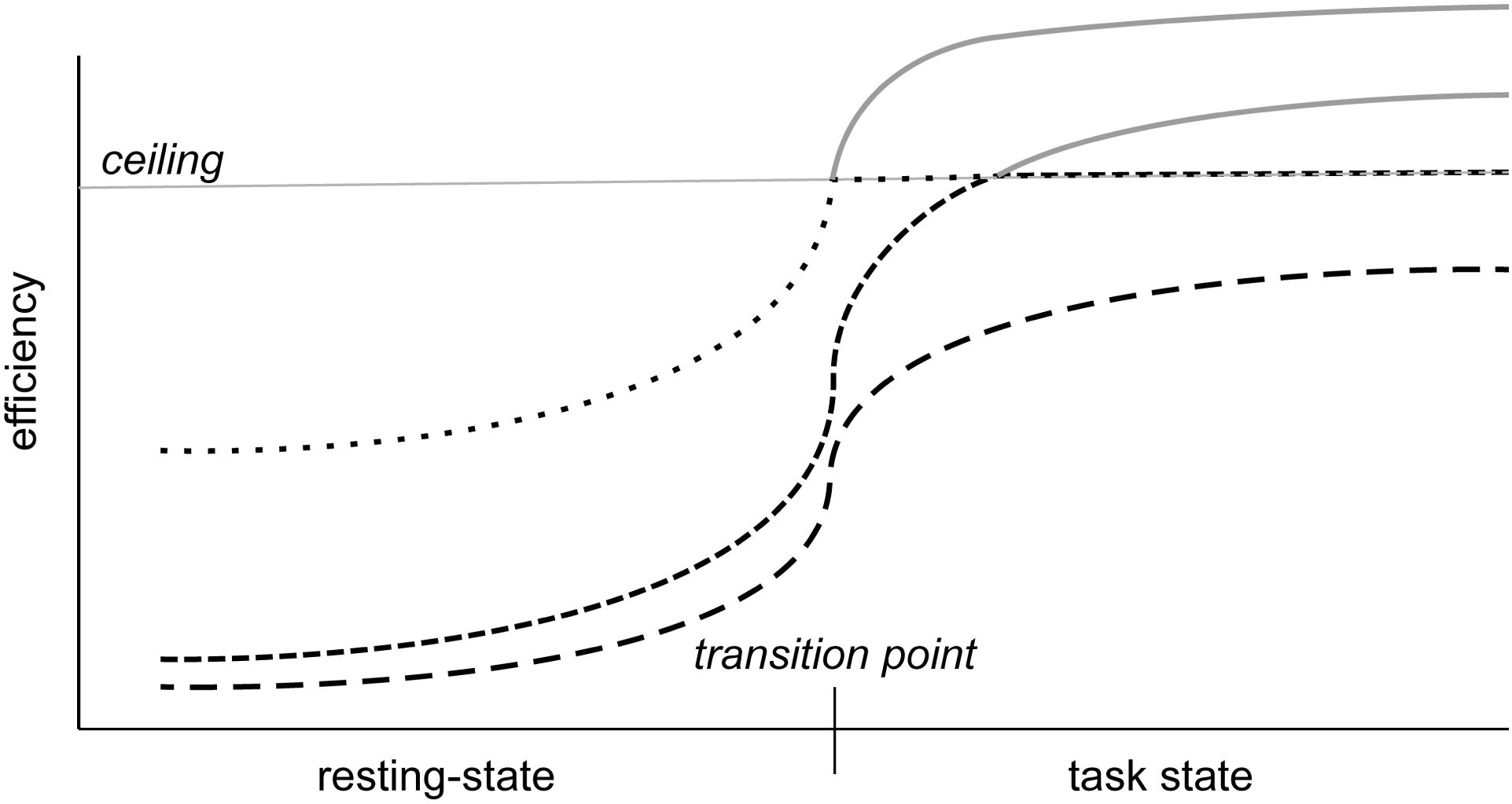
Schematic representation of rest-to-task reconfiguration hypothesis. The figure shows three fictional subjects that transition from a resting-state to task state and show a concomitant increase in (global) efficiency. The top two subjects, already have such a high efficiency during resting-state that when the brain network needs to reconfigure to a more integrated state to meet task demands, efficiency cannot surpass the ceiling (horizontal dotted lines) and leads to slower responses.

At the subnetwork level, we showed that our global results were mainly driven by inter-individual differences in FC of the DMN; both the topology of the DMN and FC between the DMN and the other RSNs (mainly the FPN) were associated with slower task performance. Because closer inspection showed that efficiency and clustering of the DMN were highly correlated (r = 0.84), the observed positive associations should instead be interpreted as an association between slower performance and increased *within* DMN FC. Indeed, when looking at total FC within the DMN, we observed a positive association (β = .25, P_bca_ = .02) with ToL reaction time. It is generally accepted that activity within the DMN is high when a subject is not engaged in any specific task and its activity is suppressed when external stimuli demand cognitive engagement (Anticevic et al. 2012). Heightened DMN activity and higher FC between the DMN and other RSNs are also commonly associated with reduced cognitive performance in brain disorder-related deficits (Putcha et al. 2016; Esposito et al. 2018; Anticevic et al. 2012). Our associations between slower ToL performance and increased within DMN FC and increased connectivity between the DMN and the other RSNs is therefore in line with these findings and adds to the growing body of literature that shows that inter-individual differences in FC of the DMN is associated with cognitive performance, even in normally functioning healthy subjects. It must be noted that these associations did not survive the multiple comparison correction, although the reported associations between performance speed at higher task load and within DMN FC and FC between DMN and FPN in our post-hoc analysis did pass the FDR correction.

In recent years, scientific awareness has increased for the low reproducibility of neuroimaging findings (Nichols et al. 2017). Test-retest reliability is often used as a measure for reproducibility and generalizability. Although graph measures, such as global efficiency and Gcc, show fair-to-good test-retest reliability (Welton et al. 2015), a recent meta-analysis showed that edges within a functional connectivity matrix – on the basis of which graph measures are calculated – show poor test-retest reliability (Noble et al. 2019). This low test-retest reliability influences statistical power and necessitates the inclusion of larger samples to reach the effect size of interest (Matheson 2019; Zuo et al. 2019). It is, however, important to note that test-retest reliability is not the same as validity and the meta-analysis showed that one of the main factors that influenced test-retest reliability was artefact correction (Noble et al. 2019); a necessary step during preprocessing to remove motion and other non-neural physiological noise from the data and avoid spurious results (Parkes et al. 2018). Moreover, although absolute values of intra-individual edges show low reproducibility (Noble et al. 2019), inter-individual differences in the functional connectome are stable across, days, months and even years (Horien et al. 2019; Finn et al. 2015; Miranda-Dominguez et al. 2014), and its characteristics are uniquely associated with a particular individual across time (Horien et al. 2019). This provides justification for predicting a person’s phenotype, including cognitive functioning, on the basis of between-subject variability of the functional connectome. Another part of reproducibility is transparent and complete reporting of the methods and results. To that end we report the COBIDAS checklist in the supplementary material (Nichols et al. 2017). We will make the here reported data available to researchers upon reasonable request.

A limitation of this study is that we exclusively looked at resting-state FC to predict performance on the ToL and not –at task-based FC, i.e. during execution of the ToL itself. This would have allowed us to look directly at the network characteristics associated with performance and to test our hypothesis of reduced ability to network integration when transitioning from rest to task. Furthermore, although conscious state may alter network topology, we did not include an objective measure to ensure wakefulness during the eyes-closed resting-state scan. A strength of this study is that we retrospectively recruited a relatively large number of healthy subjects and used stringent control for (micro)motion by excluding subjects with >0.2 mm mean RMS displacement, denoising rs-fMRI for motion-related artifacts with ICA-AROMA, employing wavelet coherence to construct the connectivity matrices and adding RMS displacement to the regression model.

In conclusion, we showed that higher global efficiency during rest and higher FC of the DMN with other RSNs and within itself is associated with slower planning performance. We tentatively postulate that due to ceiling effects individuals with a higher integrative network state during rest are less able to reconfigure to a more integrated state during task execution, leading to slower exchange across the brain network and slower behavioral responses.

## Supporting information

Cobidas

## Acknowledgements

The authors want to acknowledge Dr. Niels Gerrits and Dr. Stella de Wit for carrying out the data collection.

