## supplemental for "Resting-state network topology and planning ability in healthy adults"

**Contents**

- Supplemental Table 1 – BNA Brain Regions per subnetwork
- Supplemental Table 2 – associations between TOL reaction time and subnetwork measures
- Supplemental Table 3 – associations between TOL accuracy and subnetwork measures
- supplemental Table 4 – associations between TOL reaction time and subnetwork connectivity
- supplemental Table 5 – associations between TOL accuracy and subnetwork connectivity
- COBIDAS checklist (as separate Excel file)

| **Supplementary Table 1** - BNA Brain Regions per subnetwork | | | |
| --- | --- | --- | --- |
| BNA Node | Gyrus | BNA Label | Subnetwork |
| 175/176 | Cingulate Gyrus | A23d, dorsal area 23 | DMN |
| 179/180 | Cingulate Gyrus | A32p, pregenual area 32 | DMN |
| 181/182 | Cingulate Gyrus | A23v, ventral area 23 | DMN |
| 187/188 | Cingulate Gyrus | A32sg, subgenual area 32 | DMN |
| 33/34 | Inferior Frontal Gyrus | A45c, caudal area 45 | DMN |
| 35/36 | Inferior Frontal Gyrus | A45r, rostral area 45 | DMN |
| 39/40 | Inferior Frontal Gyrus | A44v, ventral area 44 | DMN |
| 165/166 | Insula | vIa, ventral agranular insula | DMN |
| 141/142 | Inferior Parietal Lobule | A40c, caudal area 40(PFm) | DMN |
| 143/144 | Inferior Parietal Lobule | A39rv, rostroventral area 39(PGa) | DMN |
| **[95]**/96 | Inferior Temporal Gyrus | A20il, intermediate lateral area 20 | DMN |
| 23/24 | Middle Frontal Gyrus | A8vl, ventrolateral area 8 | DMN |
| 27/28 | Middle Frontal Gyrus | A10l, lateral area10 | DMN |
| **[81]**/**[82]** | Middle Temporal Gyrus | A21c, caudal area 21 | DMN |
| **[83]**/84 | Middle Temporal Gyrus | A21r, rostral area 21 | DMN |
| 87/88 | Middle Temporal Gyrus | aSTS, anterior superior temporal sulcus | DMN |
| 41/42 | Orbital Frontal Gyrus | A14m, medial area 14 | DMN |
| 43/44 | Orbital Frontal Gyrus | A12/47o, orbital area 12/47 | DMN |
| 51/52 | Orbital Frontal Gyrus | A12/47l, lateral area 12/47 | DMN |
| 151/152 | Precuneus | dmPOS, dorsomedial parietooccipital sulcus(PEr) | DMN |
| 153/154 | Precuneus | A31, area 31 (Lc1) | DMN |
| 113/114 | Parahippocampal Gyrus | TL, area TL (lateral PPHC, posterior parahippocampal gyrus) | DMN |
| 121/122 | Posterior Superior Temporal Sulcus | rpSTS, rostroposterior superior temporal sulcus | DMN |
| 123/124 | Posterior Superior Temporal Sulcus | cpSTS, caudoposterior superior temporal sulcus | DMN |
| 3/4 | Superior Frontal Gyrus | A8dl, dorsolateral area 8 | DMN |
| 5/6 | Superior Frontal Gyrus | A9l, lateral area 9 | DMN |
| 11/12 | Superior Frontal Gyrus | A9m,medial area 9 | DMN |
| 13/14 | Superior Frontal Gyrus | A10m, medial area 10 | DMN |
| 79/80 | Superior Temporal Gyrus | A22r, rostral area 22 | DMN |
| 177/178 | Cingulate Gyrus | A24rv, rostroventral area 24 | FPN |
| 29/30 | Inferior Frontal Gyrus | A44d,dorsal area 44 | FPN |
| 31/32 | Inferior Frontal Gyrus | IFS, inferior frontal sulcus | FPN |
| 137/138 | Inferior Parietal Lobule | A39rd, rostrodorsal area 39(Hip3) | FPN |
| **[99]**/**[100]** | Inferior Temporal Gyrus | A20cl, caudolateral of area 20 | FPN |
| 17/18 | Middle Frontal Gyrus | IFJ, inferior frontal junction | FPN |
| 19/20 | Middle Frontal Gyrus | A46, area 46 | FPN |
| 21/22 | Middle Frontal Gyrus | A9/46v, ventral area 9/46 | FPN |
| 147/148 | Precuneus | A7m, medial area 7(PEp) | FPN |
| 183/184 | Cingulate Gyrus | A24cd, caudodorsal area 24 | VAN |
| 185/186 | Cingulate Gyrus | A23c, caudal area 23 | VAN |
| 107/108 | Fusiform Gyrus | A37lv, lateroventral area37 | DAN |
| 37/38 | Inferior Frontal Gyrus | A44op, opercular area 44 | VAN |
| 167/168 | Insula | dIa, dorsal agranular insula | VAN |
| 169/170 | Insula | vId/vIg, ventral dysgranular and granular insula | VAN |
| 173/174 | Insula | dId, dorsal dysgranular insula | VAN |
| 139/140 | Inferior Parietal Lobule | A40rd, rostrodorsal area 40(PFt) | DAN |
| 145/146 | Inferior Parietal Lobule | A40rv, rostroventral area 40(PFop) | VAN |
| 91/**[92]** | Inferior Temporal Gyrus | A37elv, extreme lateroventral area37 | DAN |
| 97/98 | Inferior Temporal Gyrus | A37vl, ventrolateral area 37 | DAN |
| 25/26 | Middle Frontal Gyrus | A6vl, ventrolateral area 6 | DAN |
| 15/16 | Middle Frontal Gyrus | A9/46d, dorsal area 9/46 | VAN |
| 85/86 | Middle Temporal Gyrus | A37dl, dorsolateral area37 | DAN |
| 149/150 | Precuneus | A5m, medial area 5(PEm) | DAN |
| 55/56 | Precentral Gyrus | A6cdl, caudal dorsolateral area 6 | DAN |
| 63/64 | Precentral Gyrus | A6cvl, caudal ventrolateral area 6 | DAN |
| 61/62 | Precentral Gyrus | A4tl, area 4(tongue and larynx region) | VAN |
| 7/8 | Superior Frontal Gyrus | A6dl, dorsolateral area 6 | DAN |
| 1/2 | Superior Frontal Gyrus | A8m, medial area 8 | VAN |
| 125/126 | Superior Parietal Lobule | A7r, rostral area 7 | DAN |
| 127/128 | Superior Parietal Lobule | A7c, caudal area 7 | DAN |
| 129/130 | Superior Parietal Lobule | A5l, lateral area 5 | DAN |
| 133/134 | Superior Parietal Lobule | A7ip, intraparietal area 7(hIP3) | DAN |

All odd numbers represent regions in the left hemisphere; even numbers regions in the right hemisphere. The **[##]** regions represent the regions that were excluded because they were not available across all subjects. Abbreviations: DMN = default-mode network, FPN = frontoparietal network, DAN = dorsal attention network, VAN = ventral attention network, BNA = Brainnetome Atlas. For further abbreviations of the atlas regions see: <https://atlas.brainnetome.org/bnatlas.html>

| Supplemental Table 2 – associations between TOL reaction time and subnetwork measures | | | | | | |
| --- | --- | --- | --- | --- | --- | --- |
| TOL | Model | B (SE) | 95% CI (BCa) | Beta | P_bca_ | R^2^ |
| RT | Age | 95.24 (15.7) | 61.5, 126.3 | .633 | <.001 |  |
|  | DMN eff. | 10526.9 (4323.0) | 2840.3, 21273.0 | .253 | .018 | .306 |
|  | Motion | -16036.7 (6908.2 | -30857.2, -4681.7 | -.224 | .015 |  |
|  | Age | 89.6 (16.6) | 55.2, 120.8 | .595 | <.001 |  |
|  | FPN eff | 5529.6 (4260.6) | -3457.5, 15153.3 | ..144 | .191 | .265 |
|  | Motion | -15657.5 (7437.8) | -31259.9, -2615.6 | -.219 | .033 |  |
|  | Age | 85.0 (15.7) | 50.9, 115.0 | .564 | <.001 |  |
|  | DAN eff | 2070.8 (3780.8) | -5147.6, 10618.1 | .064 | .592 | .249 |
|  | Motion | -14203.2 (6922.0) | -28020.6, -2599.0 | -.198 | .033 |  |
|  | Age | 93.7 (14.8) | 63.7, 121.8 | .623 | <.00 |  |
|  | VAN eff. | 7933.9 (4470.2) | -1231.4, 18587.6 | .214 | .079 | .287 |
|  | Motion | -18330.8 (6707.7) | -32109.9, -7779.2 | -.256 | .005 |  |
|  | Age | 90.0 (15.2) | 57.6, 120.3 | .597 | <.001 |  |
|  | DMN CC | 10489.0 (4952.2) | 2139.5, 23408.4 | .230 | .039 | .298 |
|  | Motion | -16351.0 (6506.0) | -30216.4, -5369.9 | -.228 | .050 |  |
|  | Age | 88.4 (14.6) | 59.0, 115.5 | .587 | <.001 |  |
|  | FPN CC | 8897.0 (4522.2) | 457.1, 19933,3 | .198 | .077 | .285 |
|  | Motion | -16146.7 (6698.8) | -29146.9, -5559.5 | -.225 | .055 |  |
|  | Age | 88.1 (15.0) | 56.4, 116.4 | .585 | <.001 |  |
|  | DAN CC | 8506.4 (4564.4) | 16.0, 18689.4 | .195 | .061 | .283 |
|  | Motion | -15809.5 (6550.6) | -28949.6, -5382.5 | -.221 | .013 |  |
|  | Age | 90.0 (14.9) | 56.9, 117.7 | .596 | <.001 |  |
|  | VAN CC | 9433.7 (4732.7) | 686.6, 20753.4 | .210 | .065 | .288 |
|  | Motion | -17235.9 (6412.6) | -29138.6, -7738.5 | -.241 | .043 |  |

For each analysis, age was entered in model 1, the network measure in model 2 and motion parameters in model 3. Only the results of model 3 are shown here. P-values are bootstrapped using 2000 permutations. Abbreviations: TOL = Tower of London task, RT = reaction time, ACC = accuracy, SE = Standard Error, CI = confidence interval, BCa= Bias corrected and accelerated, DMN = default-mode network, FPN = fronto-parietal network, DAN = dorsal attention network, VAN = ventral attention network. Eff = efficiency, CC = clustering coefficient. Motion was defined as the mean root-mean-squared framewise displacement during the entire resting-state MRI scan.

| Supplemental Table 3 – associations between TOL accuracy and subnetwork measures | | | | | | |
| --- | --- | --- | --- | --- | --- | --- |
| TOL | Model | B (SE) | 95% CI (BCa) | Beta | P | R^2^ |
| ACC | Age | -0.15 (0.08) | -0.32, -0.01 | -.267 | .071 |  |
|  | DMN eff. | -29.8 (24.2) | -80.3, 23.8 | -.199 | .227 | .061 |
|  | Motion | -23.7 (28.0) | -70.0, 33.4 | -.092 | .392 |  |
|  | Age | -0.14 (0.08) | -0.30, -0.003 | -.249 | .094 |  |
|  | FPN eff. | -21.7 (17.5) | -59.5, 12.6 | -.157 | .219 | .047 |
|  | Motion | -22.8 (29.6 | -73.4, 46.4 | -.089 | .406 |  |
|  | Age | -0.12 (0.07) | -0.28, 0.03 | -.217 | .129 |  |
|  | DAN eff. | -9.2 (14.1) | -34.9, 15.3 | -.079 | .528 | .030 |
|  | Motion | -28.4 (27.7) | -76.1, 36.9 | -.110 | .289 |  |
|  | Age | -0.12 (0.07) | -0.28, -0.01 | -.227 | .107 |  |
|  | VAN eff. | -8.4 (17.0) | -39.5, 18.6 | -.063 | .621 | .027 |
|  | Motion | -25.1 (28.6) | -78.3, 48.6 | -.098 | .350 |  |
|  | Age | -0.13 (0.07) | -0.30, -0.002 | -.245 | .081 |  |
|  | DMN CC | -35.1 (23.2) | -84.2, 9.8 | -.215 | .128 | .070 |
|  | Motion | -21.5 (27.9) | -68.8, 40.8 | -.084 | .422 |  |
|  | Age | -0.13 (0.08) | -0.27, 0.002 | -.236 | .098 |  |
|  | FPN CC | -30.0 (22.1) | -76.4, 11.8 | -.186 | .192 | .058 |
|  | Motion | -22.2 (29.8) | -72.2, 50.3 | -.086 | .436 |  |
|  | Age | -0.13 (0.08) | -0.286, 0.005 | -.231 | .104 |  |
|  | DAN CC | -15.1 (20.2) | -66.7, 13.7 | -.158 | .205 | .049 |
|  | Motion | -24.2 (28.6) | -72.7, 41.0 | -.094 | .376 |  |
|  | Age | -0.13 (.08) | -0.30, 0.02 | -.236 | .114 |  |
|  | VAN CC | -24.8 (21.5) | -70.7, 12.6 | -.153 | .251 | .046 |
|  | Motion | -21.0 (29.9) | -71.3, -53.0 | .081 | .444 |  |

For each analysis, age was entered in model 1, the network measure in model 2 and motion parameters in model 3. Only the results of model 3 are shown here. P-values are bootstrapped using 2000 permutations. Abbreviations: TOL = Tower of London task, RT = reaction time, ACC = accuracy, SE = Standard Error, CI = confidence interval, BCa= Bias corrected and accelerated, DMN = default-mode network, FPN = fronto-parietal network, DAN = dorsal attention network, VAN = ventral attention network. Eff = efficiency, CC = clustering coefficient. Motion was defined as the mean root-mean-squared framewise displacement during the entire resting-state MRI scan.

| supplemental Table 4 – associations between TOL reaction time and subnetwork connectivity | | | | | | |
| --- | --- | --- | --- | --- | --- | --- |
| TOL | Model | B (SE) | 95% CI (BCa) | Beta | P | R^2^ |
| RT | Age | 90.2 (15.7) | 55.3, 120.1 | .599 | <.001 |  |
|  | DMN-FPN | 9579.5 (4665.2) | 1519.4, 20298.9 | .231 | .039 | .298 |
|  | Motion | -16821.6 (6909.2) | -31410.8, -4733.7 | -.235 | .014 |  |
|  | Age | 86.3 (15.2) | 52.0, 115.7 | .573 | <.001 |  |
|  | DMN-DAN | 9525.2 (4733.2) | 995.2, 19277.1 | .205 | .042 | .288 |
|  | Motion | -15334.3 (6675.6) | -28806.7, -4974.6 | -.214 | .020 |  |
|  | Age | 87.9 (15.1) | 56.2, 117.7 | .584 | <.001 |  |
|  | DMN-VAN | 8566.3 (4350.8) | 655.4, 17947.4 | .200 | .044 | .285 |
|  | Motion | -17158.1 (6405.8) | -29479.2, -6933.1 | -.239 | .007 |  |
|  | Age | 85.5 (15.4) | 52.4, 115.1 | .568 | <.001 |  |
|  | FPN-DAN | 4867.0 (4388.8) | -3428.3, 15327.6 | .118 | .256 | .259 |
|  | Motion | -14770.7 (6970.7) | -27681.9, -4743.9 | -.206 | .024 |  |
|  | Age | 87.3 (15.4) | 52.3, 116.5 | .580 | <.001 |  |
|  | FPN-VAN | 6390.7 (4215.2) | -1399.9, 15910.1 | .151 | .117 | .267 |
|  | Motion | -16623.6 (6938.9) | -30903.6, -5240.0 | -.232 | .011 |  |
|  | Age | 89.4 (14.2) | 58.8, 117.1 | .594 | <.001 |  |
|  | DAN-VAN | 7868.2 (4142.2) | -32.0, 16651.0 | .188 | .058 | .279 |
|  | Motion | -16954.7 (6892.0) | -31729.2, -5937.1 | -.237 | .008 |  |

For each analysis, age was entered in model 1, the network measure in model 2 and motion parameters in model 3. Only the results of model 3 are shown here. P-values are bootstrapped using 2000 permutations. Abbreviations: TOL = Tower of London task, RT = reaction time, ACC = accuracy, SE = Standard Error, CI = confidence interval, BCa= Bias corrected and accelerated, DMN-FPN = functional connectivity between the default-mode network and fronto-parietal network, DMN-DAN = functional connectivity between the default mode network and dorsal attention network, DMN-VAN = functional connectivity between the default mode network and ventral attention network FPN-DAN = functional connectivity between the fronto-parietal network and dorsal attention network, FPN-VAN = functional connectivity between the fronto-parietal network and ventral attention network, DAN-VAN = functional connectivity between the dorsal and ventral attention networks. Motion was defined as the mean root-mean-squared framewise displacement during the entire resting-state MRI scan.

| supplemental Table 5 – associations between TOL accuracy and subnetwork connectivity | | | | | | |
| --- | --- | --- | --- | --- | --- | --- |
| TOL | Model | B (SE) | 95% CI (BCa) | Beta | P | R^2^ |
| ACC | Age | -0.13 (0.08) | -0.30, 0.003 | -.243 | .106 |  |
|  | DMN-FPN | -28.7 (22.2) | -75.2, 19.6 | -.192 | .211 | .060 |
|  | Motion | -21.0 (28.5) | -68.1, 34.6 | -.082 | .447 |  |
|  | Age | -0.12 (0.07) | -0.28, 0.008 | -.223 | .118 |  |
|  | DMN-DAN | -30.6 (21.8) | -79.9, 13.1 | -.183 | .166 | .058 |
|  | Motion | -25.1 (28.5) | -78.8, 41.2 | -.098 | .378 |  |
|  | Age | -0.13 (0.07) | -0.28, 0.02 | -.230 | .107 |  |
|  | DMN-VAN | -25.6 (22.3) | -73.7, 12.5 | -.166 | .243 | .051 |
|  | Motion | -20.0 (30.2) | -70.8, 45.3 | -.078 | .480 |  |
|  | Age | -0.12 (0.08) | -0.28, 0.02 | -.215 | .111 |  |
|  | FPN-DAN | -11.7 (16.8) | -46.7, 22.2 | -.080 | .533 | .030 |
|  | Motion | -27.7 (27.4) | -73.6, 28.9 | -.107 | .423 |  |
|  | Age | -0.12 (0.08) | -0.29, 0.02 | -.228 | .118 |  |
|  | FPN-VAN | -20.5 (21.2) | -65.7, 15.1 | -.134 | .349 | .041 |
|  | Motion | -21.0 (30.3) | -70.0, 46.4 | -.082 | .462 |  |
|  | Age | -0.12 (0.07) | -0.28, 0.02 | -.222 | .111 |  |
|  | DAN-VAN | -10.6 (19.3) | -49.1, 22.2 | -.070 | .587 | .028 |
|  | Motion | -25.7 (28.7) | -78.8, 40.4 | -.100 | .343 |  |

For each analysis, age was entered in model 1, the network measure in model 2 and motion parameters in model 3. Only the results of model 3 are shown here. P-values are bootstrapped using 2000 permutations. Abbreviations: TOL = Tower of London task, RT = reaction time, ACC = accuracy, SE = Standard Error, CI = confidence interval, BCa= Bias corrected and accelerated, DMN-FPN = functional connectivity between the default-mode network and fronto-parietal network, DMN-DAN = functional connectivity between the default mode network and dorsal attention network, DMN-VAN = functional connectivity between the default mode network and ventral attention network FPN-DAN = functional connectivity between the fronto-parietal network and dorsal attention network, FPN-VAN = functional connectivity between the fronto-parietal network and ventral attention network, DAN-VAN = functional connectivity between the dorsal and ventral attention networks. Motion was defined as the mean root-mean-squared framewise displacement during the entire resting-state MRI scan.
